# Microbial community-based protein derived from food-processing wastewater supports substantial fishmeal replacement and survival in Pacific white shrimp (*Litopenaeus vannamei*)

**DOI:** 10.64898/2025.12.11.693825

**Authors:** Ezequiel Santillan, Poh Leong Loo, Fanny Yasumaru, Hui Xu, Ramanujam Srinivasan Vethathirri, Yan Zhou, Diana Chan Pek Sian, Stefan Wuertz

**Affiliations:** Singapore Centre for Environmental Life Sciences Engineering, Nanyang Technological University, Singapore 637551, Singapore; Aquaculture Innovation Centre, Temasek Polytechnic, Singapore 529757, Singapore; Advanced Environmental Biotechnology Centre (AEBC), Nanyang Environment & Water Research Institute (NEWRI), Nanyang Technological University, 1 Cleantech Loop, Clean Tech One, Singapore 637141, Singapore; School of Civil and Environmental Engineering, Nanyang Technological University, Singapore 639798, Singapore

**Keywords:** Pacific white shrimp, Aquaculture, Aquafeeds, Postbiotics, Single-cell protein, Fishmeal replacement, Circular bioeconomy

## Abstract

The growing demand for aquaculture feeds requires sustainable alternatives to fishmeal. Microbial community-based single-cell protein (SCP) produced from soybean-processing wastewater offers a circular approach that combines nutrient recovery with aquafeed production. We evaluated SCP as a fishmeal replacement for Pacific white shrimp (*Litopenaeus vannamei*) across four feeding trials lasting 42–84 days, involving shrimp from different hatcheries and SCP produced at laboratory and pilot scales. Diets containing 0–100% fishmeal replacement, corresponding to up to 220 g SCP kg^−1^ feed, were tested. Replacement of 50% was consistently supported across trials, while levels up to 90% maintained several key performance outcomes, including specific growth rate, feed conversion ratio, and survival, without significant differences from the fishmeal control. Complete replacement reduced growth in one trial, potentially reflecting the lower protein content and less favourable amino acid balance of SCP relative to fishmeal, particularly its lower lysine and methionine proportions. Shrimp tissue proximate and amino acid compositions showed no clear adverse responses to SCP inclusion in the trial in which they were assessed. Cross-trial analysis revealed a positive association between SCP replacement and survival after accounting for trial-specific baseline differences, with survival increasing by an estimated 1.53 percentage points for every 10-percentage-point increase in replacement (HC3 95% CI: 0.90–2.17; *P* < 0.001). Although the mechanisms underlying this association remain to be established, the findings support investigation of potential functional effects through targeted challenge and mechanistic studies. Overall, performance across four heterogeneous trials supports substantial fishmeal replacement by microbial community-based SCP and demonstrates its potential for practical application as a circular aquafeed ingredient.

## Introduction

Aquaculture is the fastest-growing animal protein sector and plays a critical role in global food security. However, its continued reliance on fishmeal and fish oil derived from wild-capture fisheries raises concerns over sustainability and resource constraints (FAO-UN, 2020; Tacon, Metian, and McNevin, 2022). High-value aquaculture species such as shrimp still depend heavily on fishmeal as a primary protein source, highlighting the urgent need for sustainable feed alternatives.

Single-cell proteins (SCP), which include microbial biomass from bacteria, yeast, fungi, and algae, have been widely investigated as alternative protein sources (Glencross, Huyben, and Schrama, 2020; Jones *et al*., 2020). In *Litopenaeus vannamei*, several studies demonstrated the potential of SCP to partially or fully replace fishmeal. Methanotrophic bacterial SCP (*Methylococcus capsulatus*) has been shown to sustain growth and feed efficiency up to two-thirds replacement with additional benefits for immune gene expression (Felix *et al*., 2023). Complete replacement was achieved without adverse effects in some trials (Jintasataporn *et al*., 2021). However, results vary depending on the SCP source and processing method. For instance, diets containing *Corynebacterium ammoniagenes* SCP reduced growth performance above 20% replacement (Hamidoghli *et al*., 2019), while processed SCP products demonstrated improved digestibility and amino acid availability (Nederlof, Kaushik, and Schrama, 2023). These differences underscore the importance of SCP quality, production method, and strain characteristics.

Beyond growth performance, SCP can also confer functional benefits. Studies have reported enhanced disease resistance to *Vibrio* spp. challenges in shrimp fed methanotrophic SCP (Chen *et al*., 2024; Jintasataporn *et al*., 2021) and increased resilience under ammonia stress when fed purple non-sulfur bacteria (PNSB; *Rhodopseudomonas palustris*, *Rhodobacter capsulatus*) (Alloul *et al*., 2021). PNSB biomass has also demonstrated antimicrobial properties, pigment enrichment, and immunostimulatory effects in fish and shrimp (González Cámara *et al*., 2025). Such multifunctional properties position microbial proteins not only as nutritional replacements but also as health-promoting feed additives.

Most SCP studies have focused on axenic cultures of specific microorganisms. By contrast, microbial community-based SCP has recently emerged as a promising alternative with unique advantages. Community-based approaches reduce production costs by avoiding sterilisation and inoculum preparation, and they can utilise variable wastewaters directly (Vethathirri, Santillan, and Wuertz, 2021). They also deliver consistent yields and protein quality across replicates, even under fluctuating wastewater characteristics (Vethathirri *et al*., 2023; Vethathirri *et al*., 2025). In addition, some microalgal strains can accumulate polyunsaturated fatty acids when grown in effluents from SCP bioreactors, enhancing the circular benefits and supporting their use in nutrient-enriched aquafeeds (Thi *et al*., 2026). Importantly, community-based SCP produced from soybean-processing wastewater was shown to be safe for aquafeeds, with negligible levels of pathogen-like sequences and a balanced essential amino acid profile (Vethathirri *et al*., 2026; Vethathirri *et al*., 2025). Such properties underscore its potential to complement or outperform conventional single-strain SCP in real-world applications (Santillan, Neshat, and Wuertz, 2025).

Recent reviews highlight microbial proteins as among the most scalable and sustainable fishmeal alternatives (Chen *et al*., 2024; Li *et al*., 2022). While soy protein concentrate and krill meal remain widely used in shrimp feeds, both raise sustainability concerns (Barreto *et al*., 2025). Valorising food-processing wastewater into microbial protein offers a dual solution: reducing effluent treatment costs while producing sustainable aquafeed ingredients. We recently demonstrated in Asian seabass (*Lates calcarifer*) that microbial community-based SCP produced from soybean-processing wastewater could replace 50% of dietary fishmeal without compromising growth or survival (Santillan *et al*., 2026; Santillan *et al*., 2024).

Shrimp present a particular challenge because they lack adaptive immunity and cannot be vaccinated, relying instead on innate defences and diet-derived immunostimulants (Kulkarni *et al*., 2021). This places functional feed additives at the centre of shrimp health management, and recent reviews emphasise that such additives, together with omics-based approaches, will be critical for advancing shrimp nutrition and disease resistance (Chen *et al*., 2024). Microbial community-based SCP is therefore of special interest as it may combine the nutritional role of a protein replacement with functional properties linked to beneficial taxa and their postbiotic metabolites (Santillan *et al*., 2024; Tao *et al*., 2024).

Here, we extend this work by systematically evaluating microbial community-based SCP in Pacific white shrimp (*L. vannamei*), the most widely farmed crustacean species worldwide. Four independent feeding trials were conducted using different shrimp stocks and SCP produced at two scales: laboratory scale (4-L bioreactors) for Trial 1 and pilot scale (100-L bioreactors) for Trials 2–4. The trials were also longer than many previous SCP feeding studies: Trial 1 lasted 42 days, Trials 2 and 3 lasted 84 days, and Trial 4, conducted as a validation trial, lasted 56 days. These durations enabled growth, survival, and feed performance to be evaluated over extended periods. Collectively, the four trials evaluated SCP performance across heterogeneous biological and operational settings, including different shrimp stocks, SCP batches, production scales, trial durations, and inclusion ranges. This design provides evidence of performance reproducibility across settings, although it was not intended to isolate the individual effect of production scale or any other between-trial factor. We assessed growth and survival across varying levels of fishmeal replacement and descriptively compared shrimp tissue composition, providing one of the most comprehensive evaluations to date of microbial community-based SCP as a sustainable protein source for shrimp aquaculture, while also examining its potential functional value.

## Materials and methods

### Experimental animals and culture conditions

Juvenile Pacific white shrimp (*L. vannamei*) with an average initial weight of ∼1 g were used in four independent feeding trials. Shrimp originated from different hatcheries: postlarvae (PL10) from Thailand (Trial 1), juveniles from a local shrimp farm in Singapore (Trial 2), and PL10 from Malaysia (Trials 3 and 4). All batches were screened for *Enterocytozoon hepatopenaei* (EHP), and only EHP-free shrimp were used. Shrimp were stocked at 20 individuals per tank in recirculating aquaculture systems (RAS) equipped with mechanical and biological filtration and continuous aeration, using rectangular glass aquaria (0.58 m × 0.58 m × 0.38 m, 100-L water volume). Water quality was maintained at 26–28 °C, salinity 20 ppt, dissolved oxygen (DO) >6 mg LLJ¹, and pH 7.2–7.8. Ammonia, nitrite, and nitrate concentrations were within recommended levels for shrimp culture throughout the trials. All experimental procedures involving animals were performed in accordance with relevant institutional guidelines and regulations. This study was approved by the Institutional Animal Care and Use Committee (IACUC) under protocol number 202003-155A.

### Diet preparation

Experimental diets for shrimp were formulated to replace fishmeal with microbial community-based SCP produced from soybean-processing wastewater. SCP was obtained at laboratory scale for Trial 1 and at pilot scale for Trials 2–4, then dried and ground into powder. Three basal formulations contained 0, 50, or 100% replacement of fishmeal protein with SCP (SCP0, SCP50, SCP100), and in later trials intermediate inclusion levels ranging from 10 to 90% were tested. Diets were prepared by first mixing the dry ingredients, then adding water and other liquid ingredients, extruding the dough through a Hobart meat grinder, and drying overnight at 45 °C. Diets were formulated to target approximately 40% crude protein and to meet the essential amino acid requirements of *L. vannamei*. The fishmeal-based control diet (SCP0) contained 22% fishmeal, which was progressively replaced with SCP in the experimental diets, while other ingredients were kept constant (Table 1). Diets from the SCP0, SCP50, and SCP90 groups were analysed for proximate composition (protein, lipid, ash) and amino acid profiles, with crude protein determined using the Kjeldahl method, lipids and ash by gravimetric analysis, and amino acids following AOAC protocols (AOAC, 1995) (Table 2).

**Table 1.** Formulation of the fishmeal control diet (SCP0) and two experimental diets with 50% (SCP50) and 90% (SCP90) fishmeal replacement by microbial community-based single-cell protein used in the feeding trials with Pacific white shrimp (*Litopenaeus vannamei*).

| **Ingredient** | **SCP0**<br>**(%)** | **SCP50**<br>**(%)** | **SCP90**<br>**(%)** |
| --- | --- | --- | --- |
| Squid meal | 5 | 5 | 5 |
| Fish meal | 22 | 11 | 2.2 |
| SCP | 0 | 11 | 19.8 |
| Fish hydrolysate (Aquatic) | 4 | 4 | 4 |
| Dried yeast | 3 | 3 | 3 |
| Poultry by-product meal | 10 | 10 | 10 |
| Soy lecithin | 2 | 2 | 2 |
| Soybean meal | 15 | 15 | 15 |
| Wheat flour | 35 | 35 | 35 |
| Fish oil | 2 | 2 | 2 |
| Vitamin premix | 1 | 1 | 1 |
| Mineral premix | 1 | 1 | 1 |
| **Total** | **100** | **100** | **100** |

**Table 2.** Nutritional composition (as-is basis) of the fishmeal control diet (SCP0) and two experimental diets with 50% (SCP50) and 90% (SCP90) fishmeal replacement by microbial community-based single-cell protein used in the feeding trials with Pacific white shrimp (*Litopenaeus vannamei*).

| Proximate analysis (%) | SCP0 | SCP50 | SCP90 | Method |
| --- | --- | --- | --- | --- |
| *Proximate composition (%)* |  |  |  |  |
| Protein | 44.4 | 40.9 | 37.5 | A |
| Lipid | 8.8 | 8.2 | 7.8 | B |
| Ash | 9.4 | 10.3 | 10.6 | C |
| *Essential amino acid composition (%)* |  |  |  |  |
| Histidine | 0.51 | 0.45 | 0.39 | D |
| Threonine | 0.78 | 0.72 | 0.69 | D |
| Valine | 1.56 | 1.46 | 1.41 | D |
| Methionine | 0.74 | 0.59 | 0.53 | D |
| Tryptophan | 0.40 | 0.41 | 0.39 | E |
| Phenylalanine | 1.28 | 1.20 | 1.14 | D |
| Isoleucine | 1.05 | 1.00 | 0.93 | D |

<sup>A</sup> AsureQuality Method 6503s, Kjeldahl block digestion
<sup>B</sup> AsureQuality Method 4988, Gravimetric-SBR
<sup>C</sup> AsureQuality Method 4151, Gravimetric
<sup>D</sup> Analytical Biochemistry 178 (1989) (Modified).
<sup>E</sup> AsureQuality Method 5707, HPLC.
Values are based on a single pooled feed sample per dietary treatment (n = 1).

### Feeding trial design

Shrimp were fed to apparent satiation twice daily. Each diet was tested in triplicate tanks. Trial 1 aimed to evaluate the effects of partial and complete replacement of fishmeal with SCP on the performance of juvenile shrimp. The trial lasted 42 days and included SCP0, SCP50, and SCP100 diets, with endpoints including growth, survival, feed conversion ratio (FCR), and feed intake. Based on the outcomes of Trial 1, Trial 2 aimed to optimise the inclusion levels of SCP in shrimp diets, and the trial lasted 84 days with SCP inclusion levels at 0, 10, 20, 30, 40, and 50%, focusing on growth, FCR, feed intake, and survival. Trial 3 was conducted for 84 days using feeds stored for six months, with inclusion levels at 0, 10, 30, 50, 80, and 90%, and endpoints including growth, survival, FCR, and feed intake. This trial aimed to evaluate the effectiveness of 6-month-old experimental feeds on shrimp performance and its validity after storage. Trial 4 lasted 56 days and tested SCP0, SCP50, SCP80, and SCP90 diets to further validate Trial 3, using the same zootechnical parameters for consistency and comparison.

### Growth performance and survival assessments

Shrimp were individually weighed at the start and end of each trial. Growth performance was evaluated by final body weight, mean weight gain (mean final weight – mean initial weight), percent weight gain (mean weight gain * mean initial weight^−1^ *100), specific growth rate (SGR = (ln_mean final weight_ – ln_mean initial weight_) * trial duration^−1^ *100), feed intake, FCR = mean feed intake * mean weight gain^−1^, and survival rate.

### Shrimp tissue composition analysis

Following the evaluation of shrimp performance across four trials, shrimp from the SCP0 and SCP90 groups sampled from Trial 4 were analysed for proximate composition (protein, lipid, ash) and amino acid profiles. Shrimp from the SCP90 treatment were selected to represent the highest fishmeal-replacement level evaluated in Trial 4. Crude protein was determined using the Kjeldahl method, lipid and ash by gravimetric analysis, and amino acids following AOAC protocols (AOAC, 1995).

### SCP production

For Trial 1, microbial community-based SCP was produced in four 4-L sequencing batch reactors operated at SCELSE on 12-h cycles with intermittent aeration for 136 days, as described previously (Santillan *et al*., 2024). Wastewater from a soybean processing company (Mr Bean Group Ltd, Singapore) served as the sole feedstock. Reactors were maintained at 30 °C and mixed at 375 rpm. Each cycle consisted of a 5–10 min feeding phase, a 180 min anoxic/anaerobic phase, a 540 min aerobic phase, and a 60 min settling phase, after which 1.35 L of supernatant was withdrawn and replaced with fresh wastewater. During the aerobic phase, dissolved oxygen (DO) was controlled between 0.2 and 0.5 mg LLJ¹, and pH remained between 6.0 and 8.5. Each reactor was equipped with pH and DO probes (Mettler Toledo), peristaltic pumps, and a water-jacketed temperature control system. For Trials 2–4, SCP was produced at pilot scale in a 100-L aerobic sequencing batch reactor constructed at NEWRI and operated for over 500 days. The reactor was inoculated with biomass from a lab-scale enrichment and fed weekly with 200–300 L of soybean-soaking wastewater obtained from the processing company and stored at 0–2 °C before use. The influent had a chemical oxygen demand (COD) of 2,140–6,800 mg LLJ¹ and low nitrogen content (5–20 mg LLJ¹ NHLJLJ-N; average C/N ≈ 100:1). External nitrogen sources (*e.g.*, inorganic nitrogen fertilizers) were added as needed to overcome nitrogen limitation. The system was operated at hydraulic retention times (HRTs) of 1–3 days with DO maintained at 0.5–1.5 mg LLJ¹. In later stages, biomass concentration was stabilised using standard membrane-based methods. SCP was harvested by sedimentation or membrane concentration, collected every 1–3 days, spray-dried, and milled prior to diet formulation.

### Microbial characterisation of microbial community-based SCP

Twelve samples of 0.5 g each were collected randomly from 2-L bottles containing 300–500 g of mixed-culture SCP to be used in the different shrimp trials. These were subjected to DNA extraction as previously described (Santillan *et al*., 2019). Amplicon preparation was done by polymerase chain reaction (PCR) using the primer set 341f/785r targeting the V3–V4 variable regions of the 16S rRNA gene (Thijs *et al*., 2017), as described in Santillan and Wuertz (2022). Bacterial 16S rRNA amplicon sequencing was done in two steps as described in Santillan *et al*. (2020). The libraries were sequenced on an Illumina MiSeq platform (v.3) with 20% PhiX spike-in and at a read-length of 300 bp paired-end. Sequenced sample libraries were processed following the DADA2 bioinformatics pipeline (Callahan *et al*., 2016). DADA2 allows inference of exact amplicon sequence variants (ASVs) providing several benefits over traditional clustering methods (Callahan, McMurdie, and Holmes, 2017). Illumina sequencing adaptors and PCR primers were trimmed prior to quality filtering. Sequences were truncated after 280 and 255 nucleotides for forward and reverse reads, respectively, the length at which average quality dropped below a Phred score of 20. After truncation, reads with expected error rates higher than 3 and 5 for forward and reverse reads, respectively, were removed. After filtering, error rate learning, ASV inference and denoising, reads were merged with a minimum overlap of 20 bp. Chimeric sequences (0.4% on average) were identified and removed. For a total of 12 samples, on average 36,151 reads were kept per sample after processing, representing 88% of the average input reads. Taxonomy was assigned using the SILVA database (v.138.2) (Glöckner *et al*., 2017).

### Statistical analysis

Univariate testing through Welch’s analysis of variance (ANOVA) with Games-Howell post hoc grouping was done using *rstatix* (Kassambara, 2020) (v.0.6.0) in R (v. 4.4.1). Tank was treated as the experimental unit, and results are reported as mean ± SD for triplicate tanks. Where multiple pairwise comparisons were performed, *P*-values were adjusted using the Benjamini–Hochberg procedure with a false-discovery rate of 5% (Benjamini & Hochberg, 1995). To evaluate the cross-trial association between SCP replacement and shrimp survival, tank-level survival percentages were analysed in Python (v.3.12.13) using a linear model with SCP replacement as a continuous predictor and trial identity as a fixed categorical effect. The model allowed each trial to have a different intercept, accounting for differences in baseline survival, while estimating one common linear slope for SCP replacement across all four trials; accordingly, the trial-specific fitted lines are parallel. Heteroscedasticity-consistent HC3 standard errors were used to calculate the 95% confidence interval and *P*-value for the common slope. HC3 provides inference that is less sensitive to unequal residual variance and adjusts each squared residual according to its leverage in the fitted model (MacKinnon & White, 1985). A secondary model including an SCP replacement × trial identity interaction was fitted to assess whether the slope differed among trials. Model calculations and statistical inference were implemented using *NumPy* (v.2.3.5) (Harris *et al*., 2020), *pandas* (v.2.2.3) (McKinney, 2010), and *SciPy* (Virtanen *et al*., 2020). Individual tank observations and treatment means were plotted at their exact SCP replacement levels using *Matplotlib* (v.3.10.8) (Hunter, 2007), with transparency used to indicate overlapping observations. Heat maps for bacterial relative abundances of the SCP were constructed using the *DivComAnalyses* package (v.0.9) in R (Constancias & Sneha-Sundar, 2022).

## Results and Discussion

### SCP supported shrimp growth up to high inclusion levels

Across four feeding trials, shrimp fed diets containing up to 90% SCP achieved growth that was generally comparable to fishmeal controls (SCP0), although responses varied among trials and endpoints. In Trial 1, shrimp fed SCP100 had significantly lower weight gain and specific growth rate (SGR) than those fed SCP0, while shrimp fed SCP50 showed intermediate values. In Trial 2, no significant differences in growth performance were detected among treatments up to 50% replacement. In Trial 3, shrimp fed SCP90 performed comparably to those fed SCP0 in terms of total weight gain, SGR, and FCR, although some intermediate levels showed modest reductions. In Trial 4, individual weight gain was higher in SCP0 than in SCP50–90, but SGR did not differ significantly among treatments (Table 3). Although the diets were formulated to target comparable protein contents, compositional analysis showed that measured protein content generally declined as SCP replacement increased. The maintenance of several performance endpoints at high replacement despite this decline also suggests that improved protein and amino acid balancing could further enhance performance.

**Table 3.**
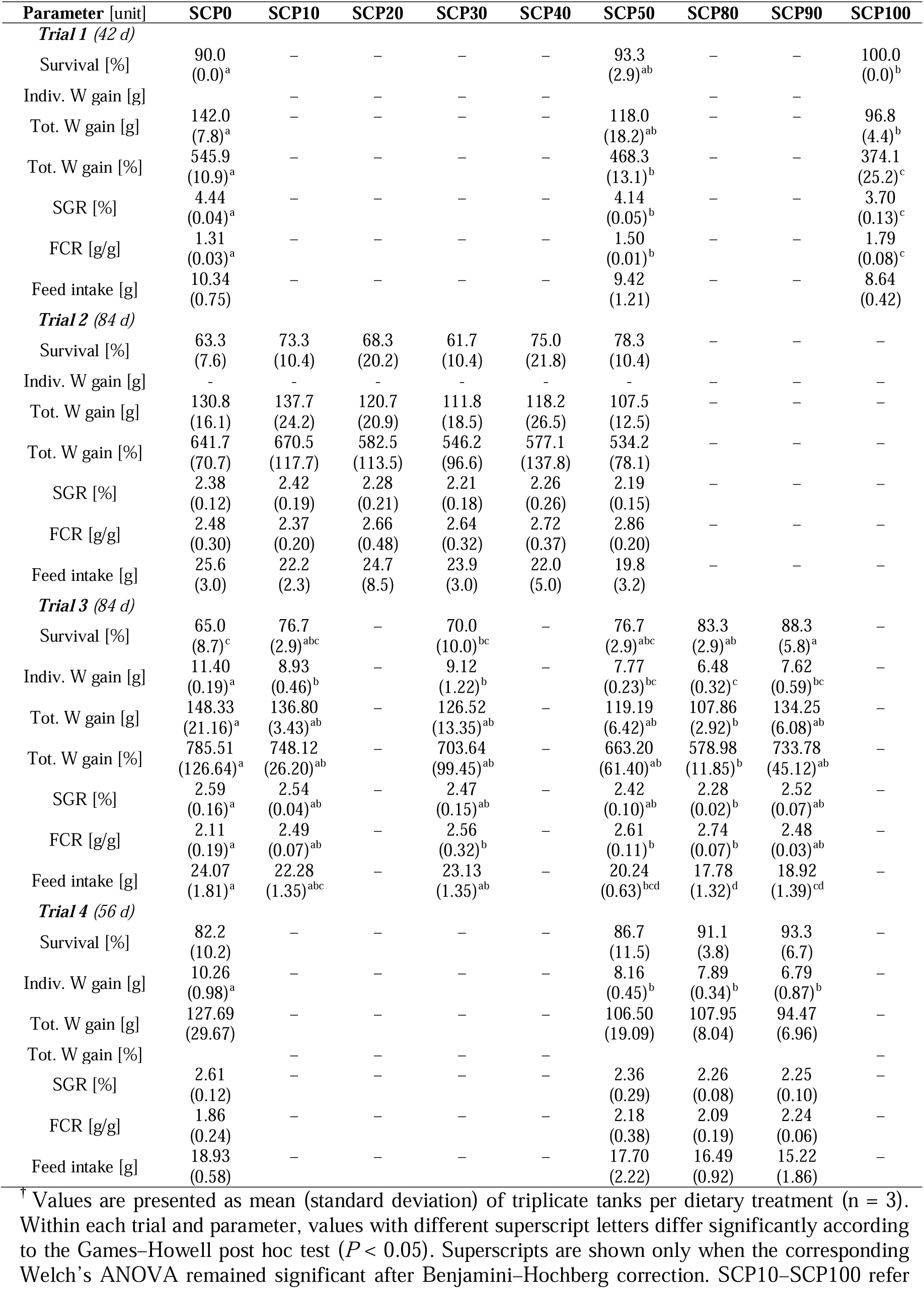
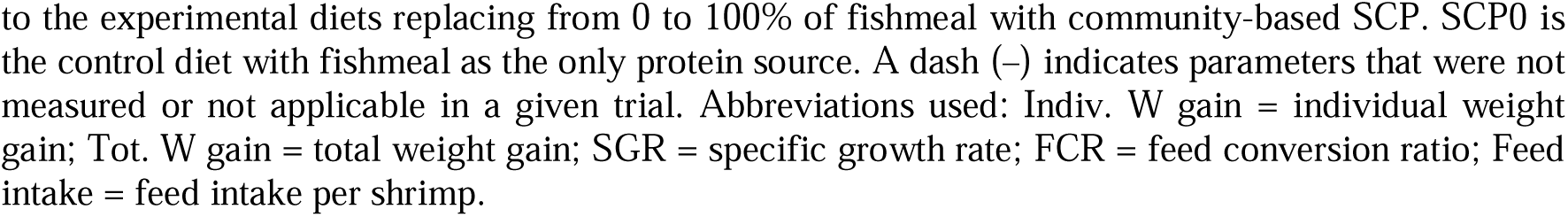
Growth performance of shrimp (*Litopenaeus vannamei*) fed diets with microbial community-based SCP across four trials^†^.

| Parameter [unit] | SCP0 | SCP10 | SCP20 | SCP30 | SCP40 | SCP50 | SCP80 | SCP90 | SCP100 |
| --- | --- | --- | --- | --- | --- | --- | --- | --- | --- |
| **Trial 1 (42 d)** |  |  |  |  |  |  |  |  |  |
| Survival [%] | 90.0<br>(0.0)<sup>a</sup> | — | — | — | — | 93.3<br>(2.9)<sup>ab</sup> | — | — | 100.0<br>(0.0)<sup>b</sup> |
| Indiv. W gain [g] | — | — | — | — | — | — | — | — | — |
| Tot. W gain [g] | 142.0<br>(7.8)<sup>a</sup> | — | — | — | — | 118.0<br>(18.2)<sup>ab</sup> | — | — | 96.8<br>(4.4)<sup>b</sup> |
| Tot. W gain [%] | 545.9<br>(10.9)<sup>a</sup> | — | — | — | — | 468.3<br>(13.1)<sup>b</sup> | — | — | 374.1<br>(25.2)<sup>c</sup> |
| SGR [%] | 4.44<br>(0.04)<sup>a</sup> | — | — | — | — | 4.14<br>(0.05)<sup>b</sup> | — | — | 3.70<br>(0.13)<sup>c</sup> |
| FCR [g/g] | 1.31<br>(0.03)<sup>a</sup> | — | — | — | — | 1.50<br>(0.01)<sup>b</sup> | — | — | 1.79<br>(0.08)<sup>c</sup> |
| Feed intake [g] | 10.34<br>(0.75) | — | — | — | — | 9.42<br>(1.21) | — | — | 8.64<br>(0.42) |
| **Trial 2 (84 d)** |  |  |  |  |  |  |  |  |  |
| Survival [%] | 63.3<br>(7.6) | 73.3<br>(10.4) | 68.3<br>(20.2) | 61.7<br>(10.4) | 75.0<br>(21.8) | 78.3<br>(10.4) | — | — | — |
| Indiv. W gain [g] | — | — | — | — | — | — | — | — | — |
| Tot. W gain [g] | 130.8<br>(16.1) | 137.7<br>(24.2) | 120.7<br>(20.9) | 111.8<br>(18.5) | 118.2<br>(26.5) | 107.5<br>(12.5) | — | — | — |
| Tot. W gain [%] | 641.7<br>(70.7) | 670.5<br>(117.7) | 582.5<br>(113.5) | 546.2<br>(96.6) | 577.1<br>(137.8) | 534.2<br>(78.1) | — | — | — |
| SGR [%] | 2.38<br>(0.12) | 2.42<br>(0.19) | 2.28<br>(0.21) | 2.21<br>(0.18) | 2.26<br>(0.26) | 2.19<br>(0.15) | — | — | — |
| FCR [g/g] | 2.48<br>(0.30) | 2.37<br>(0.20) | 2.66<br>(0.48) | 2.64<br>(0.32) | 2.72<br>(0.37) | 2.86<br>(0.20) | — | — | — |
| Feed intake [g] | 25.6<br>(3.0) | 22.2<br>(2.3) | 24.7<br>(8.5) | 23.9<br>(3.0) | 22.0<br>(5.0) | 19.8<br>(3.2) | — | — | — |
| **Trial 3 (84 d)** |  |  |  |  |  |  |  |  |  |
| Survival [%] | 65.0<br>(8.7)<sup>c</sup> | 76.7<br>(2.9)<sup>abc</sup> | — | 70.0<br>(10.0)<sup>bc</sup> | — | 76.7<br>(2.9)<sup>abc</sup> | 83.3<br>(2.9)<sup>ab</sup> | 88.3<br>(5.8)<sup>a</sup> | — |
| Indiv. W gain [g] | 11.40<br>(0.19)<sup>a</sup> | 8.93<br>(0.46)<sup>b</sup> | — | 9.12<br>(1.22)<sup>b</sup> | — | 7.77<br>(0.23)<sup>bc</sup> | 6.48<br>(0.32)<sup>c</sup> | 7.62<br>(0.59)<sup>bc</sup> | — |
| Tot. W gain [g] | 148.33<br>(21.16)<sup>a</sup> | 136.80<br>(3.43)<sup>ab</sup> | — | 126.52<br>(13.35)<sup>ab</sup> | — | 119.19<br>(6.42)<sup>ab</sup> | 107.86<br>(2.92)<sup>b</sup> | 134.25<br>(6.08)<sup>ab</sup> | — |
| Tot. W gain [%] | 785.51<br>(126.64)<sup>a</sup> | 748.12<br>(26.20)<sup>ab</sup> | — | 703.64<br>(99.45)<sup>ab</sup> | — | 663.20<br>(61.40)<sup>ab</sup> | 578.98<br>(11.85)<sup>b</sup> | 733.78<br>(45.12)<sup>ab</sup> | — |
| SGR [%] | 2.59<br>(0.16)<sup>a</sup> | 2.54<br>(0.04)<sup>ab</sup> | — | 2.47<br>(0.15)<sup>ab</sup> | — | 2.42<br>(0.10)<sup>ab</sup> | 2.28<br>(0.02)<sup>b</sup> | 2.52<br>(0.07)<sup>ab</sup> | — |
| FCR [g/g] | 2.11<br>(0.19)<sup>a</sup> | 2.49<br>(0.07)<sup>ab</sup> | — | 2.56<br>(0.32)<sup>b</sup> | — | 2.61<br>(0.11)<sup>b</sup> | 2.74<br>(0.07)<sup>b</sup> | 2.48<br>(0.03)<sup>ab</sup> | — |
| Feed intake [g] | 24.07<br>(1.81)<sup>a</sup> | 22.28<br>(1.35)<sup>abc</sup> | — | 23.13<br>(1.35)<sup>ab</sup> | — | 20.24<br>(0.63)<sup>bcd</sup> | 17.78<br>(1.32)<sup>d</sup> | 18.92<br>(1.39)<sup>cd</sup> | — |
| **Trial 4 (56 d)** |  |  |  |  |  |  |  |  |  |
| Survival [%] | 82.2<br>(10.2) | — | — | — | — | 86.7<br>(11.5) | 91.1<br>(3.8) | 93.3<br>(6.7) | — |
| Indiv. W gain [g] | 10.26<br>(0.98)<sup>a</sup> | — | — | — | — | 8.16<br>(0.45)<sup>b</sup> | 7.89<br>(0.34)<sup>b</sup> | 6.79<br>(0.87)<sup>b</sup> | — |
| Tot. W gain [g] | 127.69<br>(29.67) | — | — | — | — | 106.50<br>(19.09) | 107.95<br>(8.04) | 94.47<br>(6.96) | — |
| Tot. W gain [%] | — | — | — | — | — | — | — | — | — |
| SGR [%] | 2.61<br>(0.12) | — | — | — | — | 2.36<br>(0.29) | 2.26<br>(0.08) | 2.25<br>(0.10) | — |
| FCR [g/g] | 1.86<br>(0.24) | — | — | — | — | 2.18<br>(0.38) | 2.09<br>(0.19) | 2.24<br>(0.06) | — |
| Feed intake [g] | 18.93<br>(0.58) | — | — | — | — | 17.70<br>(2.22) | 16.49<br>(0.92) | 15.22<br>(1.86) | — |

<sup>†</sup> Values are presented as mean (standard deviation) of triplicate tanks per dietary treatment (n = 3). Within each trial and parameter, values with different superscript letters differ significantly according to the Games–Howell post hoc test ( $P < 0.05$ ). Superscripts are shown only when the corresponding Welch’s ANOVA remained significant after Benjamini–Hochberg correction. SCP10–SCP100 refer
to the experimental diets replacing from 0 to 100% of fishmeal with community-based SCP. SCP0 is the control diet with fishmeal as the only protein source. A dash (–) indicates parameters that were not measured or not applicable in a given trial. Abbreviations used: Indiv. W gain = individual weight gain; Tot. W gain = total weight gain; SGR = specific growth rate; FCR = feed conversion ratio; Feed intake = feed intake per shrimp.

Collectively, these findings indicate that SCP can replace a substantial proportion of fishmeal in shrimp diets, with replacement levels up to 90% supporting several key growth outcomes, although not all growth responses were maintained consistently at high inclusion across trials. The SCP biomass provided substantial protein and a generally favourable essential amino acid profile, but contained less protein overall and lower proportions of lysine and methionine than fishmeal (Table 4). Consequently, the reduced growth observed at complete replacement in Trial 1 may partly reflect the lower protein content and less favourable amino acid balance of the high-SCP diet. It could also be influenced by the known limitations of microbial proteins rich in nucleic acids (Felix *et al*., 2023). Bacterial SCP typically contain 8–12% nucleic acids, predominantly RNA (Ritala *et al*., 2017). Although excessive nucleic acid intake can reduce digestibility, post-treatment can substantially reduce RNA content (Vethathirri *et al*., 2026). However, nucleic acids are not exclusively detrimental: in shrimp, moderate dietary RNA has been associated with enhanced immune responses and protection against infection (Rairat *et al*., 2022). Thus, their influence on shrimp performance may depend on both the inclusion level and the broader dietary context. Importantly, SCP supported substantial fishmeal replacement across trials involving different shrimp stocks, SCP batches, production scales, trial durations, and inclusion ranges, providing evidence of performance across heterogeneous experimental settings. Comparable findings have been reported in bacterial SCP studies, where processing methods improved digestibility and amino acid availability (Nederlof, Kaushik, and Schrama, 2023). This aligns with previous studies using bacterial SCP, such as *Methylococcus capsulatus*, which supported high replacement levels in *L. vannamei* (Felix *et al*., 2023; Jintasataporn *et al*., 2021). By comparison, heterotrophic SCPs such as *Corynebacterium ammoniagenes* performed poorly above moderate inclusions (Hamidoghli *et al*., 2019), underscoring the nutritional robustness of microbial community-based SCP.

**Table 4.** Total amino acid-derived protein content and essential amino acid composition of fishmeal and microbial community-based SCP used in whiteleg shrimp feeding trials^†^.

| Component | Fishmeal | SCP |
| --- | --- | --- |
| Total amino acid-derived protein (%) | 55.1 (2.0) | 40.0 (1.5) |
| Arginine (% protein) | 5.97 (0.16) | 5.42 (0.16) |
| Histidine (% protein) | 2.28 (0.06) | 2.06 (0.74) |
| Isoleucine (% protein) | 3.23 (0.11) | 3.34 (0.54) |
| Leucine (% protein) | 7.62 (0.23) | 7.70 (0.40) |
| Lysine (% protein) | 10.19 (0.31) | 4.72 (0.63) |
| Methionine (% protein) | 2.83 (0.12) | 2.01 (0.32) |
| Phenylalanine (% protein) | 3.79 (0.17) | 4.15 (0.44) |
| Threonine (% protein) | 4.87 (0.13) | 5.61 (0.23) |
| Valine (% protein) | 4.84 (0.16) | 5.05 (0.56) |

<sup>†</sup> Values are presented as mean (standard deviation). Fishmeal values were determined from triplicate samples (n = 3), whereas SCP values were determined from six independent samples collected across Trials 1 and 2 (n = 6). Essential amino acid concentrations were normalized to total amino acid-derived protein and are expressed as a percentage of protein.

### High SCP inclusion maintained and sometimes enhanced survival

Survival remained high across treatments in Trials 1 and 4, with treatment means ranging from 90.0% to 100% and from 82.2% to 93.3%, respectively (Table 3). Survival was lower overall in Trials 2 and 3, but no SCP treatment showed significantly lower survival than its corresponding fishmeal control. In Trial 1, survival was significantly higher in SCP100 than in SCP0 and SCP50, while in Trial 3, survival reached 88.3% in SCP90 and was significantly higher than in SCP0. No significant differences among diets were detected in Trials 2 or 4. In Trial 4, the lower mean survival and higher individual weight gain observed in SCP0 could reflect greater mortality-associated release of resources or possible cannibalism; however, mortality causes and shrimp behaviour were not directly assessed, and this interpretation therefore remains speculative.

Cross-trial analysis further supported a positive association between SCP replacement and survival (Fig. 1). After accounting for trial-specific differences in baseline survival, the model estimated a 1.53-percentage-point increase in survival for every 10-percentage-point increase in SCP replacement (HC3 95% CI: 0.90–2.17; P < 0.001). Trial-specific slopes were positive in all four experiments, and no significant SCP replacement × trial interaction was detected (*P* = 0.68), providing no evidence that the relationship differed among trials. Thus, the overall association was not driven solely by differences in baseline survival among experiments. Given the importance of survival to shrimp-production performance, the absence of adverse effects at high replacement levels, together with the positive cross-trial association, supports the potential value of microbial community-based SCP beyond its contribution as a protein source.

**Fig. 1.**
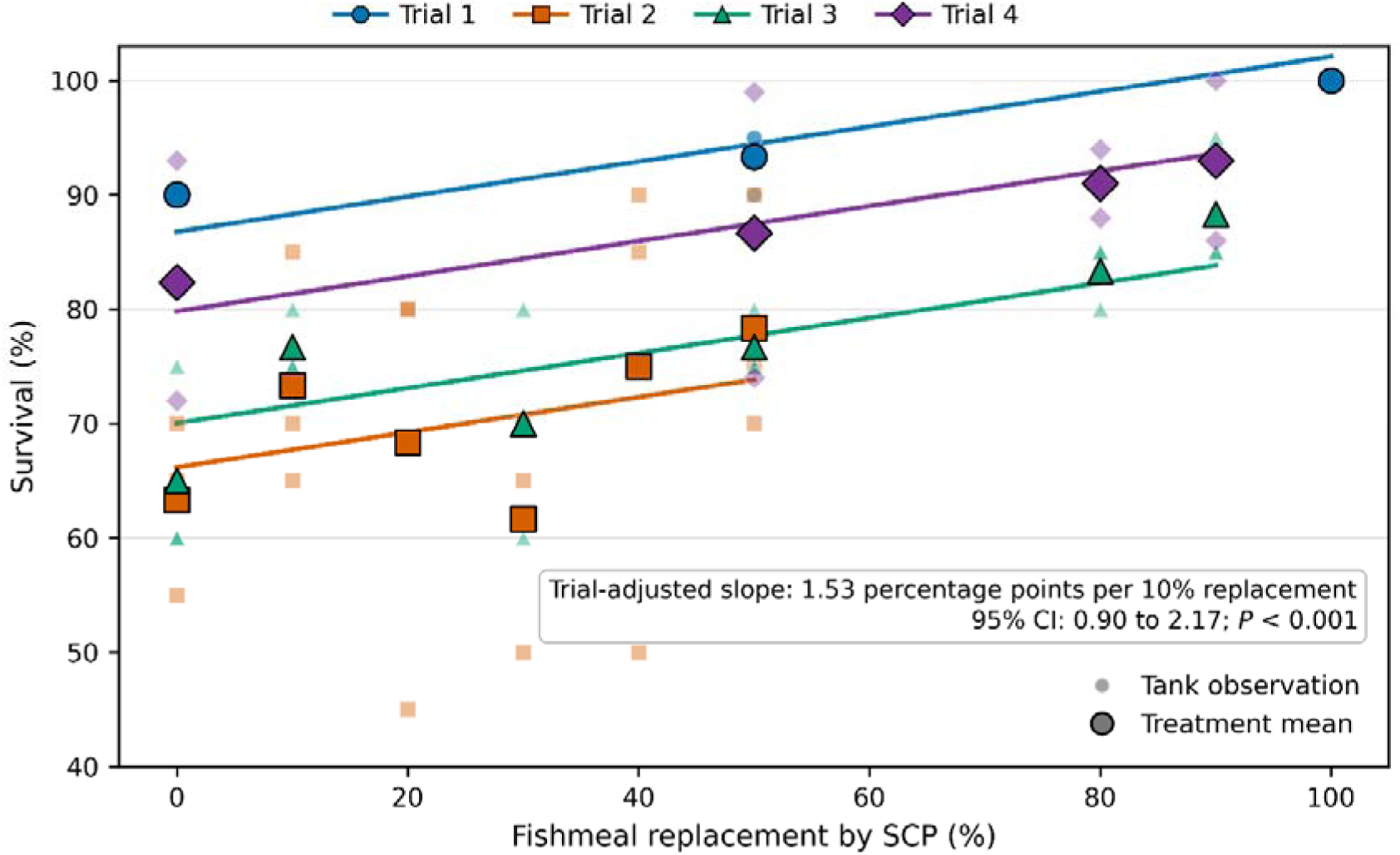
Association between fishmeal replacement by microbial community-based single-cell protein (SCP) and survival of Pacific white shrimp (*Litopenaeus vannamei*) across four independent feeding trials. Small semi-transparent symbols represent individual tank observations, while larger outlined symbols indicate treatment means. Colours and symbol shapes distinguish Trial 1 (42 days), Trial 2 (84 days), Trial 3 (84 days), and Trial 4 (56 days). Lines show predictions from a linear model with SCP replacement as a continuous predictor and trial identity as a fixed effect, allowing trial-specific intercepts while estimating a common slope. Survival increased by an estimated 1.53 percentage points for every 10-percentage-point increase in SCP replacement (HC3 95% CI: 0.90–2.17; *P* < 0.001). Tank observations are plotted at their exact SCP replacement levels, with transparency indicating overlapping observations.

Microbial cellular components and metabolites may contribute immunostimulatory or postbiotic effects, as reported for methanotrophic SCP associated with enhanced resistance to *Vibrio* infection (Chen *et al*., 2024; Jintasataporn *et al*., 2021) and for purple non-sulfur bacteria associated with improved stress tolerance and pathogen suppression (Alloul *et al*., 2021; González Cámara *et al*., 2025). However, the present trials did not include pathogen challenges, immune measurements, or mechanistic analyses, and the observed survival association cannot be attributed directly to these mechanisms. Nevertheless, its occurrence across four independent trials lasting 42–84 days, involving shrimp from different hatcheries and SCP produced at laboratory and pilot scales, demonstrates performance across substantial biological and operational variability. Targeted challenge and mechanistic studies are now needed to determine whether SCP-derived components directly contribute to shrimp resilience.

### Descriptive comparison of shrimp tissue composition

Proximate composition and amino acid profiles were measured in pooled shrimp samples from the SCP0 and SCP90 treatments in Trial 4. Because only one pooled sample was analysed per treatment, these data provide a descriptive comparison and were not subjected to inferential statistical testing. The two pooled samples had broadly similar protein, ash, and essential amino acid profiles, whereas lipid content was lower in the SCP90 sample (Table 5). These observations provide no indication of a major compositional divergence between the two samples, but replicated analyses would be required to determine whether dietary SCP affects shrimp tissue composition or nutritional value. The broadly comparable profiles are consistent with findings from our previous Asian seabass (*Lates calcarifer*) trials (Santillan *et al*., 2026; Santillan *et al*., 2024) and other SCP feeding studies in which high dietary inclusion did not significantly affect muscle composition (Felix *et al*., 2023; Jintasataporn *et al*., 2021). Nevertheless, replicated studies conducted over a complete production cycle and under commercial farming conditions are needed to determine whether these compositional patterns are maintained across populations and production settings. Such evidence would help establish whether microbial community-based SCP can reduce reliance on fishmeal while maintaining the nutritional quality of harvested shrimp, an important consideration in the development of more sustainable aquafeeds (Tacon, Metian, and McNevin, 2022).

**Table 5.** Nutritional composition (freeze-dried basis) of shrimp (*Litopenaeus vannamei*) fed control (SCP0) and high-inclusion SCP90 diets in Trial 4 over 56 days.

| Proximate analysis (%) | SCP0 | SCP90 | Method |
| --- | --- | --- | --- |
| *Proximate composition (%)* |  |  |  |
| Protein | 76.4 | 78.0 | A |
| Lipid | 6.1 | 4.6 | B |
| Ash | 11.5 | 12.6 | C |
| *Essential amino acid composition (%)* |  |  |  |
| Histidine | 1.06 | 1.06 | D |
| Threonine | 1.31 | 1.30 | D |
| Valine | 2.44 | 2.49 | D |
| Methionine | 1.40 | 1.35 | D |
| Tryptophan | 0.68 | 0.70 | E |
| Phenylalanine | 3.29 | 3.36 | D |
| Isoleucine | 1.67 | 1.70 | D |
| Leucine | 3.76 | 3.80 | D |
| Lysine | 4.03 | 4.03 | D |

<sup>A</sup>ASUREQuality Method 6503s, Kjeldahl block digestion
<sup>B</sup>ASUREQuality Method 4988, Gravimetric-SBR
<sup>C</sup>ASUREQuality Method 4151, Gravimetric
<sup>D</sup>Analytical Biochemistry 178 (1989) (Modified).
<sup>E</sup>ASUREQuality Method 5707, HPLC.
Values are descriptive and based on a single pooled shrimp sample per dietary treatment (n = 1).

### Microbial diversity and postbiotic potential position SCP as a prospective functional feed ingredient for shrimp

The microbial communities present in the SCP used in the shrimp diets were diverse and compositionally distinct across trials, reflecting changes in reactor scale, operating conditions, and SCP production batches (Fig. 2). Early SCP batches used in Trial 1 were dominated by members of the *Bacillota* phylum, whereas pilot-scale batches used in Trials 2–4 showed greater representation of *Pseudomonadota*, *Bacteroidota*, and *Actinomycetota* (Fig. 2A). Families of particular relevance included members of the *Lactobacillaceae*, *Streptococcaceae*, *Comamonadaceae*, *Prevotellaceae*, and *Pseudonocardiaceae* (Fig. 2B). Members of these taxonomic groups encompass bacteria capable of producing compounds such as organic acids, exopolysaccharides, and antimicrobial metabolites (Wegh *et al*., 2019). However, taxonomic composition alone does not establish that these compounds were produced or retained in the SCP biomass. The presence of lactic acid bacteria, including members of the *Lactobacillaceae* and *Streptococcaceae*, is nevertheless relevant to the potential functional value of the SCP. Preparations derived from *Lactobacillus*, *Lactococcus*, and *Weissella* can contain microbial cells, structural components, and metabolites associated with immune modulation, pathogen inhibition, or feed preservation, including after microbial inactivation (Quintanilla-Pineda *et al*., 2023; Tao *et al*., 2024; Thakur *et al*., 2025). *Bifidobacterium*, detected in some SCP batches, has likewise been associated with the production of organic acids and extracellular polysaccharides with potential immunomodulatory properties (Meena *et al*., 2025). Members of the *Prevotellaceae* and *Pseudonocardiaceae* are known for synthesizing polysaccharides and antimicrobial metabolites that may support gut barrier protection and stress resilience (Quintanilla-Pineda *et al*., 2023; Wegh *et al*., 2019), although their functional contributions within the present SCP remain to be established experimentally.

**Fig. 2.**
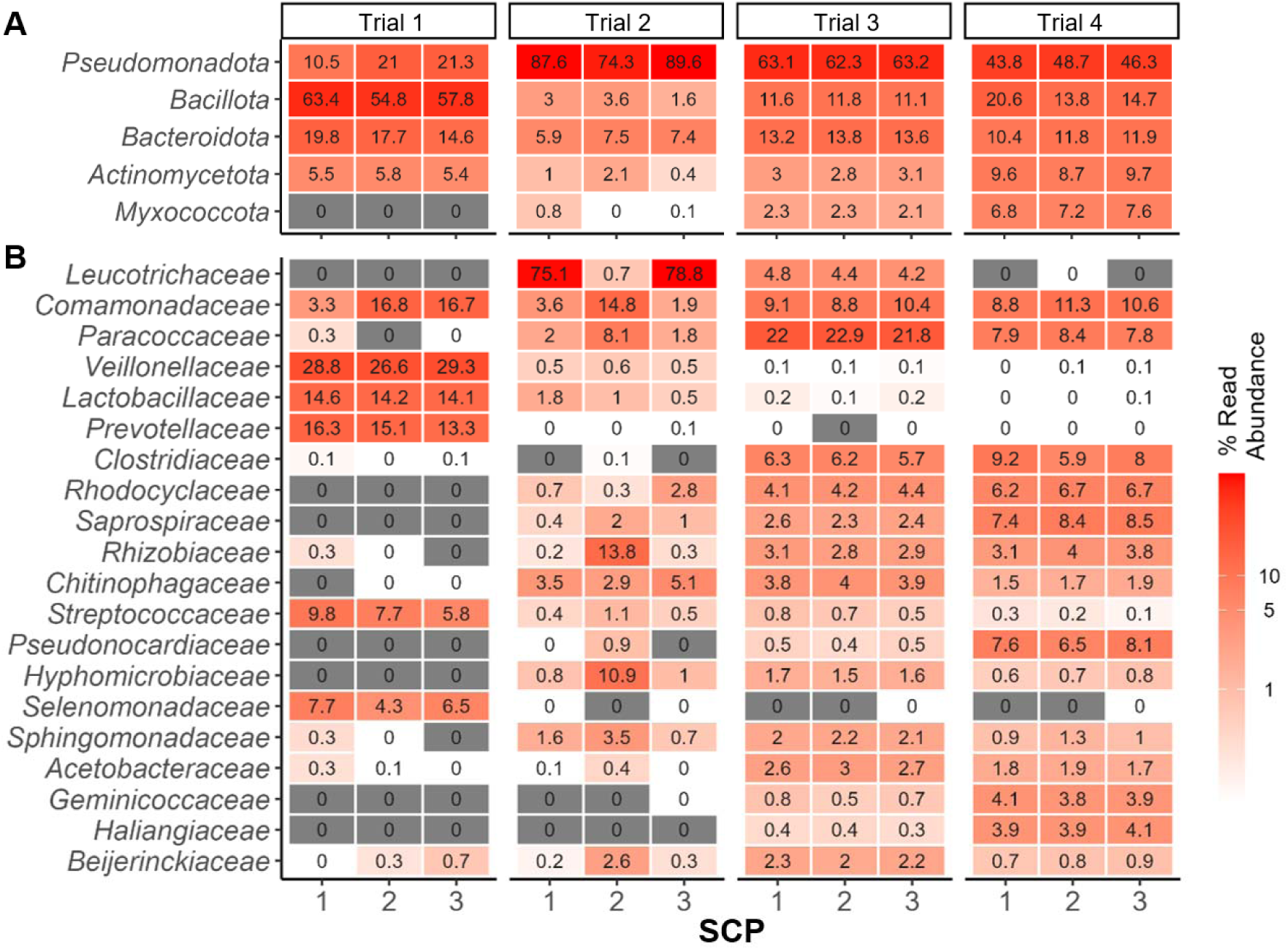
Characterisation of the microbial community-based SCP produced from soybean-processing wastewater, assessed by 16S rRNA gene amplicon sequencing. Heatmaps show the relative abundances of the (A) five most abundant phyla and (B) 20 most abundant families in triplicate SCP samples from each trial (n = 3 per trial).

Spray drying at an inlet temperature exceeding 180 °C would be expected to substantially reduce microbial viability while potentially retaining heat-stable metabolites and structural cell components. Such retained components could contribute bioactive functionality to the SCP biomass by supporting nutrient utilisation, epithelial integrity, and pathogen resistance (Thakur *et al*., 2025), although microbial viability and the retention and activity of specific compounds were not measured here. Importantly, despite variation in microbial composition across trials, substantial fishmeal replacement generally maintained shrimp performance relative to the fishmeal control, although responses at the highest replacement levels were trial- and endpoint-dependent (Table 3). This consistency across compositionally distinct SCP batches is compatible with the functional redundancy and metabolic breadth expected of microbial community-based SCP. Together, these findings suggest that community-derived SCP may offer value beyond its protein content, with potential benefits for feed functionality, shrimp health, and production robustness that warrant direct investigation. Whereas several axenic SCP products have shown reduced performance at high dietary inclusion under particular experimental conditions (Hamidoghli *et al*., 2019; Kuo *et al*., 2024; Samsing *et al*., 2024), community-based SCP offers broader metabolic diversity and potential functional redundancy, which may support process stability under non-sterile, large-scale production conditions (Vethathirri *et al*., 2026; Vethathirri *et al*., 2023; Vethathirri *et al*., 2025; Vethathirri, Santillan, and Wuertz, 2021).

A particularly promising dimension of this added value is the postbiotic potential of microbial community-based SCP. Postbiotics include bioactive compounds such as short-chain fatty acids, vitamins, bacteriocins, exopolysaccharides, and microbial cell wall components, which can exert immunomodulatory, antimicrobial, and metabolic effects (Meena *et al*., 2025; Wegh *et al*., 2019). These compounds act through mechanisms that modulate gut-associated immune responses, strengthen epithelial barriers, and inhibit pathogens. This is especially relevant for shrimp, which lack vertebrate-like adaptive immunity and therefore cannot benefit from conventional vaccination in the same manner as vertebrate aquaculture species, relying primarily on innate defences and dietary immunostimulants (Kulkarni *et al*., 2021). In the present study, survival showed a positive association with increasing SCP replacement after accounting for differences in baseline survival among trials (Fig. 1). This association occurred without an imposed pathogen or chemical challenge and cannot be attributed directly to immune stimulation or postbiotic activity. Nevertheless, it provides a compelling hypothesis for targeted investigation, particularly because increasing SCP inclusion was not associated with reduced survival even at high fishmeal-replacement levels. Related evidence supports the biological plausibility of this hypothesis: yeast-derived RNA and nucleotides have improved shrimp immune responses and survival during *Vibrio* challenge (Rairat *et al*., 2022), while PNSB biomass has increased tolerance to ammonia stress and resistance to pathogens (Alloul *et al*., 2021). In contrast, plant proteins such as soy protein concentrate have been linked to impaired immune responses in shrimp postlarvae (Barreto *et al*., 2025), highlighting the importance of alternatives that do not compromise, and may even strengthen, host defences.

Unlocking the contribution of postbiotics to shrimp performance will require direct chemical, biological, and mechanistic validation. Targeted metabolomics could quantify candidate organic acids, polysaccharides, nucleotides, vitamins, and antimicrobial compounds, while genome-resolved metagenomics and other multi-omics approaches could identify the microbial genes and pathways associated with their production (Neshat *et al*., 2024). These analyses should be coupled with controlled pathogen or environmental-stress challenges and measurements of immune, intestinal-barrier, and health-related responses. Artificial intelligence and machine-learning approaches may subsequently help integrate microbial, metabolic, process, and animal-performance data to identify taxa–function relationships and predict formulations associated with improved outcomes (Santillan, Neshat, and Wuertz, 2025). If the proposed functionality is confirmed, microbial community-based SCP could be developed not merely as a lower-value substitute for fishmeal protein, but as a differentiated feed ingredient combining circular nutrient recovery, nutritional value, and postbiotic functionality for shrimp aquaculture.

## Conclusions

Microbial community-based single-cell protein (SCP) produced from soybean-processing wastewater represents a promising alternative to fishmeal in Pacific white shrimp (*Litopenaeus vannamei*) diets. Evidence across four heterogeneous feeding trials supports substantial fishmeal replacement, with 50% emerging as a consistently supported level and replacement up to 90% maintaining several key performance outcomes, including SGR, FCR, and survival, without significant differences from the fishmeal control in the later trials. The generally favourable findings across four trials lasting 42–84 days, involving shrimp from different hatcheries and SCP produced at laboratory and pilot scales, demonstrate performance across substantial biological and operational variability and strengthen the potential for practical application. These findings support 90% replacement as a promising upper range under the tested formulations, while improved protein and amino acid balancing may further enhance performance at high inclusion and potentially enable complete replacement in diets with comparable fishmeal content. Increasing SCP inclusion did not adversely affect survival and may offer additional functional value, although the underlying mechanisms require targeted challenge and mechanistic studies. Further work should include replicated assessment of shrimp tissue composition and validation over complete production cycles under commercial farming conditions. By integrating nutrient recovery with aquafeed production, this approach offers a practical route to reduce reliance on marine-derived protein and advance more circular shrimp-production systems.

## Data availability

DNA sequencing data are available at NCBI BioProject PRJNA890376.

## Author Contributions

ES, PLL, FY, DCPS and SW conceived the studies and designed the experiments. SW and DCPS obtained the funding for the study. HX, RSV and ES produced microbial protein. PLL provided the PL10 shrimp used in Trials 3 and 4. PLL and FY performed the aquaculture trials. PLL gathered and compiled all data from aquaculture trials and associated chemical analyses. ES performed the statistical analyses, coordinated the molecular work and performed the bioinformatics analyses. ES and PLL coordinated chemical analyses. ES and PLL interpreted the data and generated the results. ES wrote the first draft of the manuscript. All authors reviewed the manuscript.

